# Metagenomic analyses reveal the influence of depth layers on marine biodiversity in tropical and subtropical regions

**DOI:** 10.1101/2022.10.18.512769

**Authors:** Bianca C. F. Santiago, Iara D. de Souza, João Vitor F. Cavalcante, Diego A. A. Morais, Rodrigo J. S. Dalmolin

## Abstract

The emergence of open ocean global-scale studies provided important information about the genomics of oceanic microbial communities. Metagenomic analyses shed a light on the structure of marine habitats, unraveling the biodiversity of different water masses. Many biological and environmental factors can contribute to marine organism composition, such as depth. However, much remains unknown about the taxonomic and functional features of microbial communities in different water layer depths. Here, we performed a metagenomic analysis of 76 samples from the Tara Ocean Project, distributed in 8 collection stations located in tropical or subtropical regions, and sampled from three layers of depth (surface water layer – SRF, deep chlorophyll maximum layer – DCM, and mesopelagic zone – MES). In total, we assigned genomic sequences to 669.713.333 organisms. The SRF and DCM depth layers are similar in abundance and diversity, while the MES layer presents greater diversity than the other layers. Diversity clustering analysis shows differences regarding the taxonomic content of samples. At the domain level, bacteria prevail in the majority of samples, and the MES layer presents the highest proportion of archaea among all samples. A core of essential biological functions was identified between the depth layers, such as DNA replication, translation, transmembrane transport, and DNA repair. However, some biological functions were found exclusively in each depth layer, suggesting different functional profiles for each of them. Taken together, our results indicate that the depth layer influences microbial sample composition and diversity.

## Introduction

Marine biodiversity consists of the variety of life present in the sea at different levels of complexity, from taxa to ecosystems (Sala et al., 2021). Even though the oceans cover 71% of the earth’s surface (Grosberg et al., 2012), only 16% of the planet’s described species are marine (Costello and Chaudhary, 2017). It is believed that most of the undescribed species are found in the marine environment (Costello et al., 2012) and the exploration of deep-sea areas can increase new species discovery (Aguzzi et al., 2019). In the last decades, open sea expeditions provided relevant amounts of ocean data, especially microbiome data. A successful example is the TaraOceans expedition (Sunagawa et al., 2015), which investigated 210 collection stations in 20 biogeographic provinces and collected samples of seawater and plankton from the Red Sea, Mediterranean Sea, Atlantic Ocean, Indic Ocean, Pacific Ocean, and Southern Ocean (Pesant et al., 2015; dev, 2015).

Marine microorganisms composition varies depending on latitude and depth (Paoli et al., 2022). TaraOceans project collected samples from three depth layers: the surface water layer (SRF - between 3 m and 7 m deep), deep chlorophyll maximum layer (DCM - between 7 m and 200 m deep), and mesopelagic zone (MES - between 200 m and 1000 m deep). The sea microbiome is complex, composed of viruses, bacteria, archaea, and unicellular eukaryotes, which are responsible for the impulse for the biogeochemistry global cycles that regulate the Earth system. While the small viruses are ubiquitous and the plankton more abundant in seawater, prokaryotes do 30% of primary production and 95% of community respiration in oceans (Pesant et al., 2015).

For a long time, microbiology studied only species that could be cultivated and analyzed in the laboratory. Plate counting for determining microbial variability is a standard method in microbiology. However, this technique is limited to research on environmental microorganisms. It has been estimated that about 1% of microorganisms are electable to be cultivated (Davey, 2011; Allen et al., 2004). Therefore, traditional culture methods are unable to detect the great majority of environmental microorganisms (Barbesti et al., 2000). With the advances in the next sequence technologies, metagenomics analysis emerged as a solution to deeply investigate complex microbial communities, such as those from marine environments (Garza and Dutilh, 2015). The metagenomic analysis does not need to obtain pure cultures for sequencing. The samples are obtained from communities and not from isolated populations, allowing hypotheses about the interactions between the communities members (Victor et al., 2008). Therefore, metagenomics allowed researchers to study the genomes of most microorganisms that cannot be easily obtained in pure culture (Hugenholtz, 2002).

Here, we investigated microbiological communities from eight different collection stations in the tropical and subtropical zones of the globe, in three different depth layers. From the data provided by TaraOceans, we selected collection stations present in the tropical and subtropical belts of the globe that presented samples in the three depth layers simultaneously. The samples were grouped, classified, and functionally annotated with the MEDUSA pipeline (Morais et al., 2022). Clustering analysis of sample diversity separates the MES layer from the others. We also found that SRF and DCM depth layers show considerable similarity in terms of composition, abundance, and diversity, while the MES layer presents a significantly greater diversity difference than the others.

## Methods

### Data selection

Data was downloaded from the Tara Oceans expedition project (Sunagawa et al., 2015). We selected 76 samples (collected between July 2010 and June 2011) from 8 tropical/subtropical collection stations. The selected stations spread across the Indian Ocean (stations 064 and 065), the South Atlantic Ocean (stations 068, 076, and 078), and the South Pacific Ocean (stations 098, 111, and 112). Each station presents data from three depth layers: (*i*) surface water layer (SRF, between 3 m and 7 m deep), (*ii*) deep chlorophyll maximum layer (DCM, between 7 m and 200 m deep) and (*iii*) mesopelagic zone (MES, between 200 m and 1000 m deep). Supplementary Table S1 shows the number of samples in each station by depth.

### Metagenomics analysis workflow

Metagenomic analysis was performed with the MEDUSA pipeline (Morais et al., 2022). Fastp tool was used to remove adapters and low-quality sequences, and to transform paired-end sequences into single-end ones (Chen et al., 2018). Fastx-collapser tool collapsed the duplicated sequences into the single-end samples resulting from the previous step (Gordon and Hannon, 2010). The resulting sequences were aligned against the non-redundant (NR) protein bank of the National Center for Biotechnology Information (NCBI) using the Diamond tool (Buchfink et al., 2015). The taxonomic classification was performed with the kaiju software (Menzel et al., 2016). This software also uses the NCBI NR database, containing all proteins belonging to Archaea, Bacteria, and Viruses, to classify the sequences in seven taxonomic ranks (domain, phylum, class, order, family, genus, and species).

### Biological diversity analysis

We investigated the microbiome biodiversity by evaluating the diversity and the abundance indices across different stations and depth layers in tropical/subtropical oceans. Diversity was estimated through the Shannon-Weaver Index (SWI) for each sample (Shannon, 1948; Shannon and Weaver, 1949). SWI is influenced by sample richness, i.e., the number of different species, as well as sample evenness, i.e., the species distribution uniformity (Kim et al., 2017). As the richness and the evenness increase in a given sample, SWI also increases. SWI was calculated as the following. Consider the *p*_*ij*_ the proportion of species *i* in sample *j*. The total number of species in sample *j* is *S*_*j*_. Therefore, the SWI for sample *j* is given by *H*_*j*_ (Equation 1):

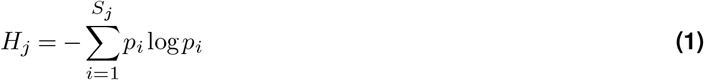

To compare SWI among the samples, SWI was normalized by the total number of species (*M*) across all samples (Equation 2).

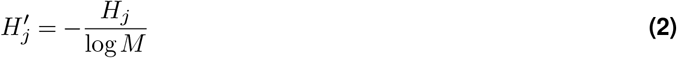

Additionally, the abundance index was estimated as the total number of organisms in a given sample.

### Clustering analysis

We performed an unsupervised clustering analysis to explore the similarity of microbial diversity between different stations and layers. The unweighted pair group method with arithmetic mean (UPGMA) was used to cluster samples based on their normalized SWI. This method calculates the euclidean distance between any pair of samples, resulting in groups of samples hierarchically similar based on their diversity profiles. All 76 samples were used for clustering analysis. The optimal number of clusters was defined by the Mojena method (Mojena, 1977).

### Functional analysis

The sequences were annotated according to the Gene Ontology (GO) database (**?**). We explored the GO biological process terms that stand out between samples in different layers. We calculated the total number of term occurrences by layer. We represented the 50 most frequent terms in each layer by a word cloud.

## Results

### Biological diversity analysis

Here, we investigate the microbiological composition of three depth layers (SRF, DCM, and MES) in eight collection stations distributed among the Indian, the South Atlantic, and the South Pacific oceans (Supplementary Figure S1). There is a noticeable heterogeneity between the samples (Figure 1). Figure 1A compares the diversity index against the abundance by sample. Sample abundance ranged from 51,666 to 39,643,916 organisms, with a median of 4,475,369 organisms. The microbiome diversity is measured by Shannon-Weaver Index (See Methods for details). The diversity index ranged from 0.40 to 0.64, with samples from the MES layer showing greater diversity than the samples from SRF and DCM layers (Figure 1A). As the majority of stations provided multiple samples, we averaged the diversity and abundance indexes by each station. The MES layer stands out showing greater diversity than the other layers (Figure 1B). Additionally, the MES layer is consistently more diverse when considering the diversity index across stations (Figure 1C). The abundance distribution is wide, with samples diverging in orders of magnitude from each other in the same station and in the same layer (Figure 1D). By layer, the MES diversity distribution differs from the SRF and DCM layers (Wilcoxon rank sum test, adjusted p-value < 0.001, Figure 2A), however, there is no significant difference in abundance distribution between the layers (Figure 2B).

**Figure 1.**
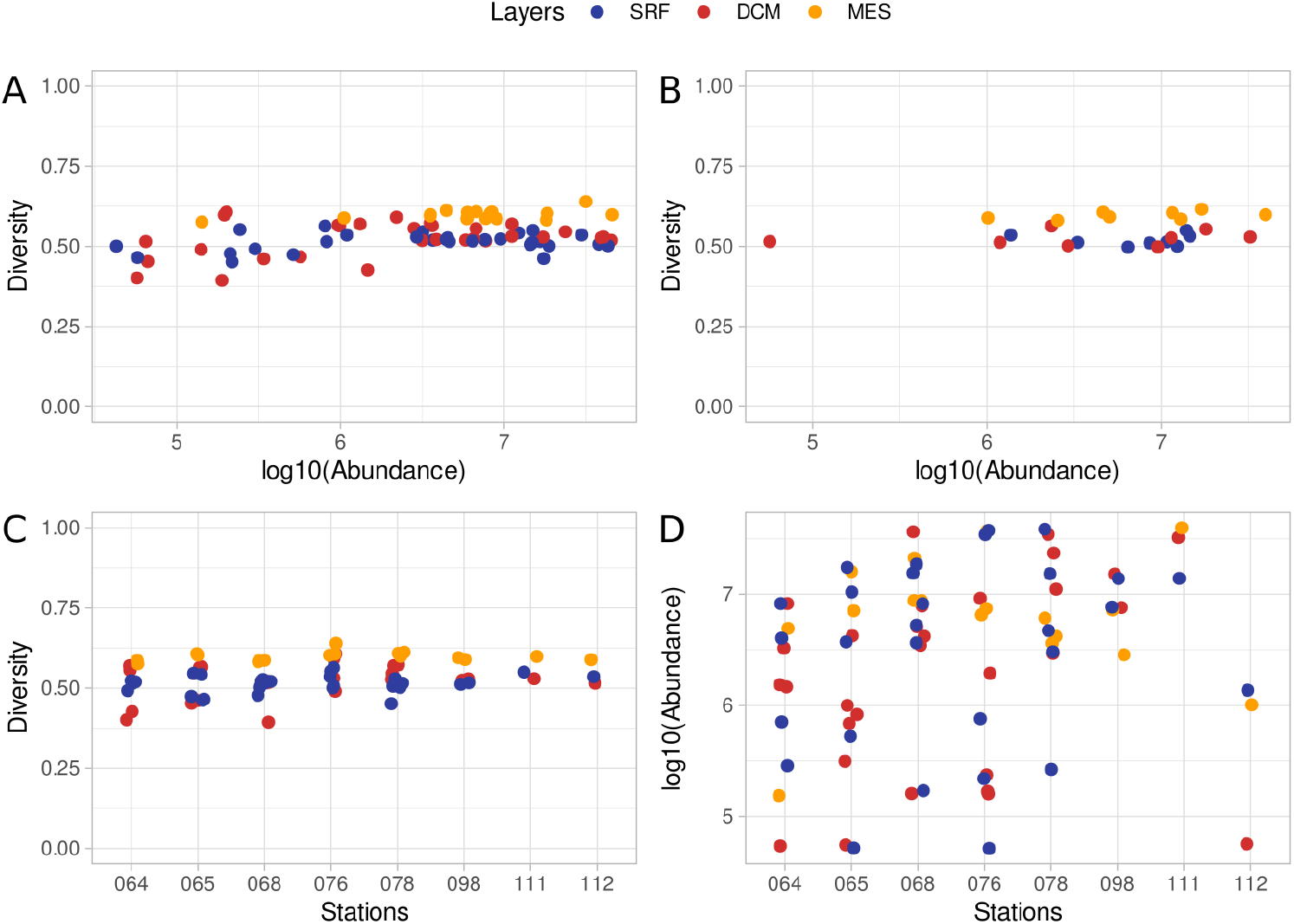
Distribution of diversity and abundance indexes. **A)** Comparison of diversity and log10-scaled abundance indexes by sample. **B)** Diversity and log10-scaled abundance averages by the station. **C)** Diversity distribution by the station. **D)** Log10-scaled abundance distribution by the station.

**Figure 2.**
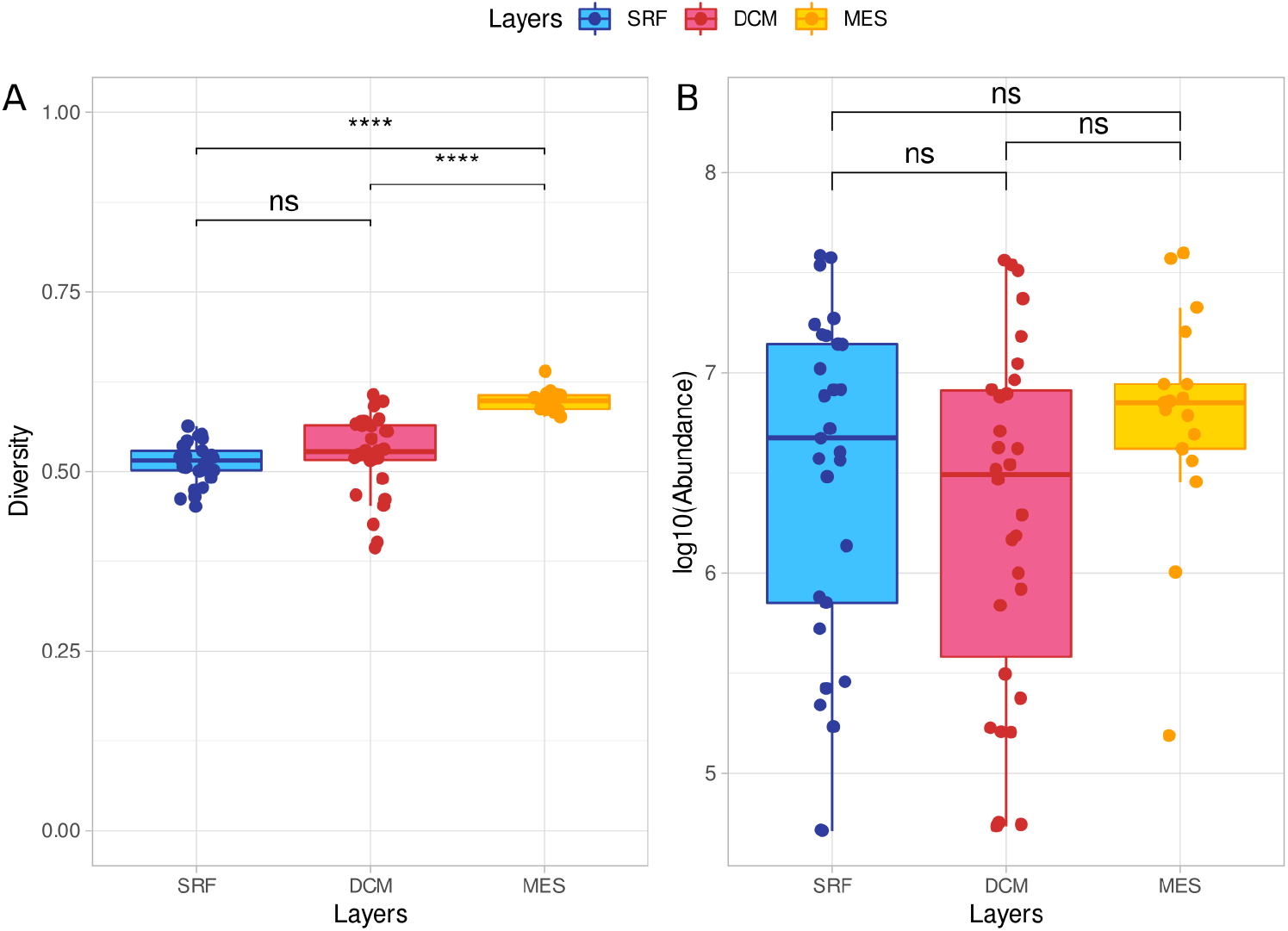
Diversity and abundance distribution by layer. **A)** Diversity comparison between layers shows increasing diversity with increasing depth. **B)** Abundance comparison between layers shows no significant association with depth layers. Comparisons were performed with the Wilcoxon rank sum test and Bonferroni-adjusted p-values were represented in each comparison. Statistical significance is represented as follows. ns (adjusted p-value > 0.05); * (adjusted p-value <= 0.05); ** (adjusted p-value <= 0.01); *** (adjusted p-value <= 0.001); **** (adjusted p-value <= 0.0001).

### Clustering and taxonomic analysis

Samples were clustered by diversity. The unsupervised analysis identified seven groups of samples, named from G1 to G7 (Figure 3A). Groups were composed of samples from different oceans, except for G1 and G5, which were composed of unique samples. This indicates that the collection location may not be decisive to sample composition. When considering the depth layers, almost all MES samples were clustered in G6, while other groups are composed of both SRF and DCM samples (Figure 3A), reinforcing our previous observation of biological diversity distinction among MES and both SRF and DCM.

**Figure 3.**
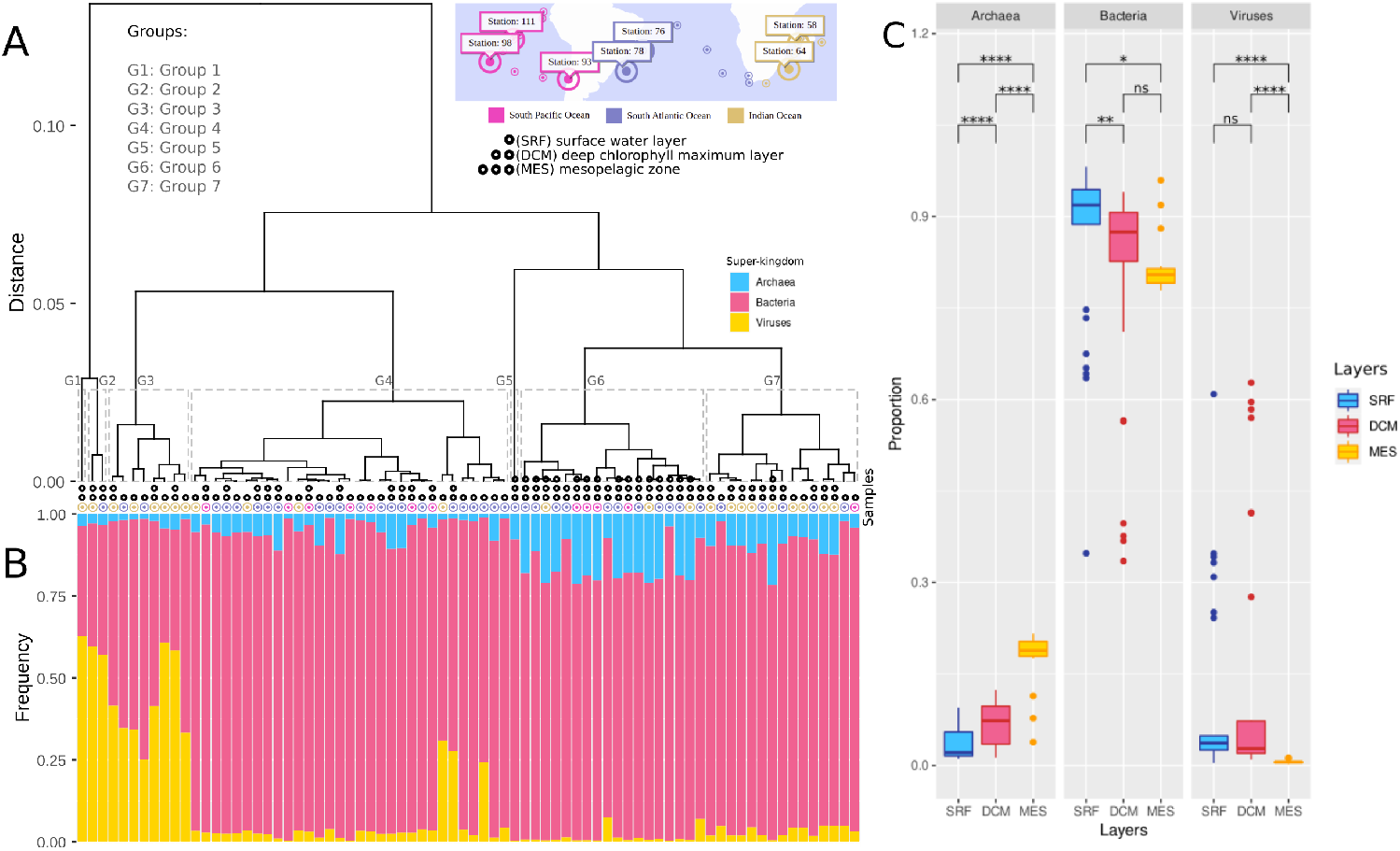
Clustering analysis of samples by diversity. **A)** Dendrogram of samples clustered by the diversity index. Seven groups of samples were identified and named from G1 to G7. Samples are also represented by depth layers and oceans. **B)** Domain-level taxonomic classification shows the proportion of each domain (archaea, bacteria, and viruses) in each sample. **C)** Distribution of domain proportions considering the samples of each depth layer. Statistical significance is represented as follows. ns (adjusted p-value > 0.05); * (adjusted p-value <= 0.05); ** (adjusted p-value <= 0.01); *** (adjusted p-value <= 0.001); **** (adjusted p-value <= 0.0001).

The taxonomic content of groups reflects the layer diversity differences. Figure 3B shows the proportion of domains (archaea, viruses, and bacteria) identified for samples in each group (Figure 3B). Bacteria prevail in groups G4 to G7. Samples from the G1 to G3 groups and a few samples from the G4 group show a high proportion of viruses. Samples from G6 show collectively the highest proportion of archaea among all groups. At the species level, the sets of overrepresented species differ between the groups (Supplementary Figure S2). Additionally, the distribution of domain proportions differs between the depth layers (Figure 3C). The proportion of archaea increases as the depth increases, with a higher proportion of archaea in DCM when compared to SRF (Wilcoxon rank sum test, adjusted p-value = 6.7E-05) and in MES when compared to DCM (Wilcoxon rank sum test, adjusted p-value = 8.2E-08). Instead, deeper layers show a decreasing proportion of bacteria (SRF to DCM, Wilcoxon rank sum test, adjusted p-value = 3.6E-03) and viruses (DCM to MES, Wilcoxon rank sum test, adjusted p-value = 2.6E-08).

### Functional content by layer

We performed an exploratory analysis of layer functional composition. We evaluated the absolute frequency of GO terms annotated for sequences. Term frequency in SRF, DCM, and MES layers are represented in Figure 4 and in Supplementary Table S2. Terms such as DNA *replication, translation, transmembrane transport, DNA repair, tricarboxylic acid cycle, carbohydrate metabolic process*, and *glutamine metabolic process* appear in the three layers. These terms represent a core of essential biological functions common to the organisms in all layers. Additionally, we evaluated terms that are present exclusively in a given layer and that represent the layer-specific functional biological profile. Twenty-four, 53, and 10 terms appear exclusively in SRF, DCM, and MES layers, respectively. We also evaluated the distribution of exclusive terms identified in each station and by each layer (Supplementary Figure S3). In spite of the greater diversity observed on the MES layer, there is no difference in the functional exclusivity between the layers. This indicates that the greater organism diversity observed in the MES layer might not be reflected in its functional diversity. The three most exclusive frequent terms in the SRF layer are *circadian rhythm, 5-phosphoribose 1-diphosphate biosynthetic process*, and *negative regulation of DNA recombination*. The three most exclusive frequent terms in the DCM layer are *heme-O biosynthetic process, intracellular protein transport*, and *phosphate-containing compound metabolic process*. Finally, the three most exclusive frequent terms in the MES layer are *methanogenesis, lysine biosynthetic process via aminoadipic acid*, and *nitrate metabolic process*.

**Figure 4.**
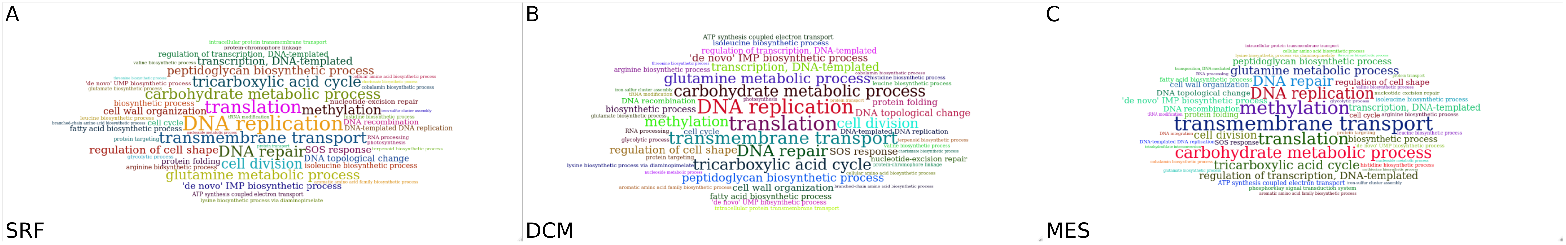
Arrangement of functional terms. **A, B, C)** Word clouds from SRF, DCM, and MES layers, respectively. The size of the terms represents their frequency in the depth layer.

## Discussion

We evaluated the organismal and functional content of subtropical marine samples in three depth layers from twelve collection stations from the Indian, the South Atlantic, and the South Pacific oceans. Abundance differences between samples from the same collection stations reflect the high sample heterogeneity. No significant difference in abundance was identified between the three depth layers studied here. However, a study showed that the vertical changes in abundance, biomass, and species composition are more accentuated than the regional differences (Auel and Hagen, 2002). Another study pointed to the sea surface temperature as the factor with the greatest influence on species abundance, showing that increases in sea surface height variation impacted the numerical abundance of euphasids (Letessier et al., 2009). The depth layers were sampled in different ways. The surface water layer was defined as the region between 3 and 7 m deep, while the deep chlorophyll maximum layer, from a chlorophyll fluorometer. The mesopelagic layer was established based on vertical profiles of temperature, salinity, fluorescence, nutrients, oxygen, and particulate matter (Pesant et al., 2015). Such a sampling strategy together with the differences in the chemical composition of the depth layers may be factors that influence the heterogeneity of the samples between the different depths.

The mesopelagic (MES) layer is more diverse than the other layers, indicating that species found in deeper layers can adapt to low temperatures and lower incidences of sunlight. A principal component analysis of marine samples showed greater differences in community composition in MES compared to both SRF and DCM samples than the differences between SRF and DCM samples (Sunagawa et al., 2015). Although the deep sea occupies 60% of the planet, the exploration of this region occurs in a much more restricted way than in coastal areas (Costello and Chaudhary, 2017). The deep sea provides an ideal scenario for species richness, given the less extreme temperatures than the ones observed in superficial layers (Grassle, 1989). A collection of heterogeneous habitats arise given the environmental conditions at bathyal depths (between 1000 m and 4000 m depth) and abyssal depths (from 4000 m depth) at different geographic and bathymetric scales (Ramirez-Llodra et al., 2011; Riehl et al., 2018). However, the level of environmental heterogeneity has unknown effects on the dispersion and distribution of species both at high depths and in inland waters (Danovaro et al., 2020). Thus, the species may have greater distribution ranges in deep waters than in shallow waters (Costello and Chaudhary, 2017).

Different patterns of marine chemical composition were observed at the different depths explored, in addition to a change in the composition of the microbial community over the months and between the different depths (cui, 2019). Bacteria are predominant in the majority of samples. Many factors contribute to the prokaryote diversity in ocean habitats, such as the availability of organic components and particulate matter, pelagic zones, and depth (Mestre et al., 2017; Ferreira et al., 2022; Walsh et al., 2016). In situ studies of deep-sea bacterioplankton showed the occurrence of bacterioplankton control by seafloor virioplankton, as there was a decrease in bacterial diversity and a change in its structure in the presence of diluted viruses (Zhang et al., 2020). Additionally, archaea assemblages were richer in deeper waters in the Arctic and Antarctic oceans, indicating that the more superficial archaea communitiesshowed lower diversity than those in the deeper layers (Xia et al., 2017).

The viral abundance showed a significant interoceanic difference for the epipelagic (0 to 200 m depth) and mesopelagic (200 to 1000 m depth) zones (Lara et al., 2022). In a metagenomic analysis of DNA virus diversity it was observed that the most abundant and widespread viral Operational Taxonomic Units (vOTUS) in South China Sea DNA virome (SCSV) are from uncultured viruses annotated from viral metagenomics, indicating that most marine viruses have not yet been characterized (Liang et al., 2019). Our results show that the three depth layers have similar relative abundances, regardless of the geographic region (collection stations), and that the MES layer is the richest in terms of diversity. In addition, despite the core of functional profiles among the organisms in the three layers, some biological functions are exclusive of each depth layer, indicating their particularities and their functional diversity.

## Code availability

Scripts used for the analyses performed here can be found at: https://github.com/dalmolingroup/meta_taraoceans (Accessed on October 17, 2022).

## Aknowledgements

The scholarships were financed by the governmental Brazilian Agency Coordination for the Improvement of Higher Education Personnel (CAPES – Portuguese: Coordenação de Aperfeiçoamento de Pessoal de Nível Superior), National Council of Technological and Scientific Development (CNPq), and PROPESQ-UFRN. We thank the STRING, SRA database, and GWAS catalog for freely providing their data. We also would like to thank NPAD/UFRN for computational resources.

## Author contributions

Conceptualization, Data curation, Formal analysis, Investigation, Methodology, Visualization, Writing– Original Draft, Writing– Review & Editing: Bianca C. F. Santiago; Data curation, Formal analysis, Investigation, Visualization, Writing– Review & Editing: Iara D. de Souza; Writing–Review & Editing: João Vitor F. Cavalcante; Data curation, Formal analysis, Investigation, Visualization: Diego A. A. Morais; Conceptualization, Supervision, Project administration, Writing– Review & Editing: Rodrigo J. S. Dalmolin.

