## Supplementary Figures for "Metagenomic analyses reveal the influence of depth layers on marine biodiversity in tropical and subtropical regions"

A

B

**Figure S1.** Latitude filter of collection stations. **A)** Filter 1: selection of stations within an interval of 5° positive or negative in relation to stations 76 and 78. **B)** Filter 2: from filter 1, selection of stations with samples in the three depth layers simultaneously.

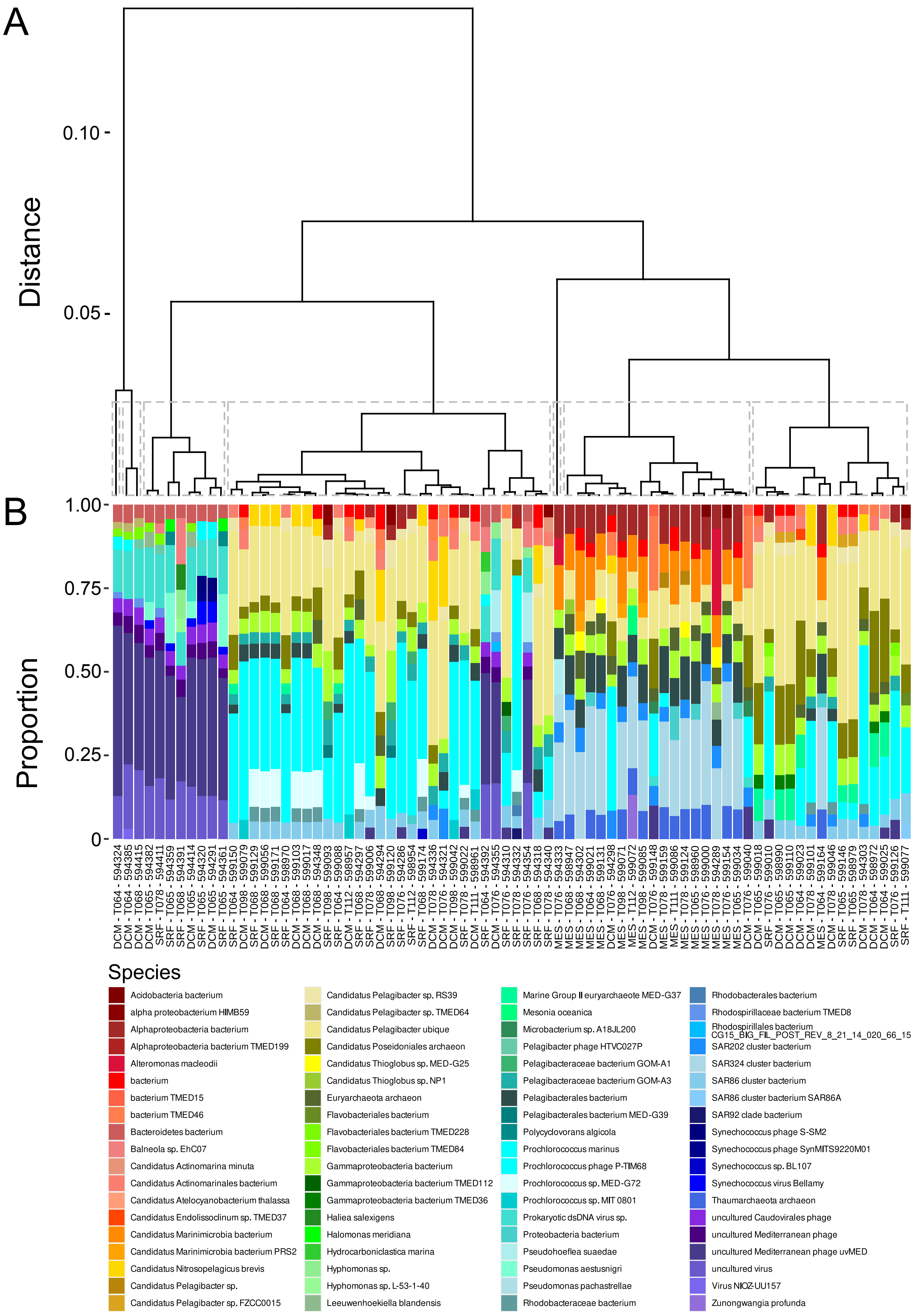


**Figure S2**. **A)** Dendrogram of samples clustered by the diversity index. **B)** Species-level taxonomic classification shows the proportion of ten most frequent species in each sample.

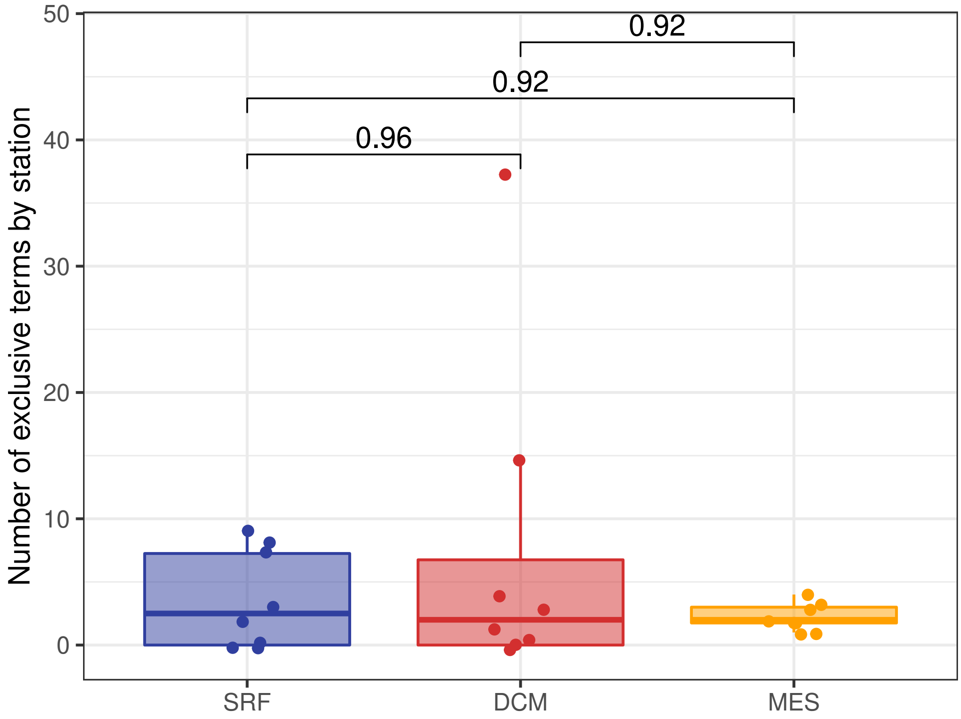


**Figure S3**: Distribution of exclusive terms by stations in each depth layer. Exclusive terms are the ones that appear only in a given depth layer of a given station. Comparisons are performed with the Wilcoxon rank sum test and Bonferroni-adjusted p-values are represented in each comparison. No significant comparison was observed.
